# The PD-L1 metabolic interactome in glioma is bounded by IDH1 genotype

**DOI:** 10.64898/2026.11.22.768411

**Authors:** Paula Carretero-Navarro, Pilar López-Larrubia, Jesús Pacheco-Torres

## Abstract

Immune checkpoint blockade has failed in glioblastoma, and part of the reason may lie in the tumour cell itself: if the checkpoint and the metabolism that sustains the cell hold each other up, then lowering one arm hands the tumour the other, and a single agent cannot win. We tested that possibility in GL261 glioma cells of IDH1-wildtype and IDH1-mutant genotype, silencing *Chka* or *Cd274*, blocking PD-L1 with an antibody, and reading the consequences by RT-qPCR, RNA sequencing, flow cytometry of surface PD-L1 and high-resolution ^1^H NMR of dual-phase cell extracts. The two genes proved reciprocal in both genotypes: silencing either transcript raised the other, and silencing choline kinase roughly doubled PD-L1 at the cell surface. Antibody blockade did something the silencing did not. In IDH1-wildtype cells it lowered soluble phosphocholine, the only one of twenty-two aqueous signals to survive correction for multiple testing, and raised total lipid per cell without changing lipid composition, whereas silencing PD-L1 raised phosphocholine instead; in IDH1-mutant cells neither change occurred. The transcriptome drew the same boundary, engaging a sterol programme and lowering glycolysis, the tricarboxylic acid cycle and oxidative phosphorylation in wild-type cells only. The interferon response did not respect that boundary and rose under every perturbation in both genotypes. Removing PD-L1 from inside the cell and blocking it from outside are therefore not the same intervention, and the metabolic half of the response stops at the IDH1 genotype.

## Introduction

Glioblastoma is the most aggressive of the diffuse gliomas of adults and remains incurable. Median survival with the standard of care is close to fifteen months and has not moved in two decades [1], so what the disease needs is not a refinement of that regimen but a treatment that works on a different principle. Immunotherapy was the obvious candidate, and it has not delivered: immune checkpoint blockade, which changed the outlook in several solid tumours, failed its phase 3 trials of PD-1 blockade in glioblastoma, both in newly diagnosed [2] and in recurrent disease [3]. Trials of that size are not defeated by sample size, and the obstacle is not access to the tumour, since blockade given before surgery does reach it and reprograms it, raising tumour-infiltrating lymphocytes and interferon-related expression [4]. Something inside the tumour restores the suppression that the antibody releases, and what that something is remains unclear.

One place to look is the metabolism of the cell that carries the checkpoint. Abnormal choline metabolism is one of the oldest and most general metabolic features of malignant transformation, and the phosphocholine that accumulates in transformed cells is the product of the first committed step of phosphatidylcholine synthesis, the Kennedy pathway [5]. The enzyme that catalyses that step, choline kinase α (ChoKα), is overexpressed in a wide range of human tumours and in tumour-derived cell lines, and its activity tracks the malignant phenotype rather than accompanying it [6]. In glioblastoma cells the phosphocholine pool responds to the signalling that sustains ChoKα [7], and inhibiting the enzyme genetically or pharmacologically suppresses their mesenchymal programme [8]. The pathway is therefore not a passive marker of the transformed state but a node the cell depends on.

PD-L1 is the ligand of the PD-1 checkpoint, and it also signals inside the tumour cell that carries it: blocking it on tumour cells lowers their mTOR activity and their glycolysis [9], and attenuating it slows their proliferation and augments their autophagy in culture [10]. Those two observations put PD-L1 and cell metabolism on the same axis, and one study has measured the axis directly. In human triple-negative breast and pancreatic cancer cell lines, ChoKα and PD-L1 behave as a reciprocal pair: silencing *CHKA* raises PD-L1 transcript and PD-L1 at the cell surface, and silencing *CD274* raises ChoKα and the phosphocholine that depends on it, so that lowering either arm is compensated by a rise in the other [11]. If that reciprocity is general, it matters for treatment, because it means that an agent directed at one arm hands the tumour the other. Whether the pair exists in glioma cells has not, to our knowledge, been examined.

Glioma is a good place to ask, because one of its defining lesions sits in a metabolic enzyme. The 2021 World Health Organization classification separates astrocytoma, IDH-mutant, CNS WHO grade 4, from glioblastoma, IDH-wildtype [12], and mutations of IDH1 and IDH2 occur in the majority of lower-grade gliomas and of the grade 4 tumours that arise from them [13]. Patients with the IDH-mutant tumour live longer, but the disease is not curable, and the one approved targeted agent lengthens progression-free survival in grade 2 disease without an overall-survival read-out [14]. Because the mutation produces 2-hydroxyglutarate, the metabolic context in which any metabolism–checkpoint relation would have to operate differs between the two genotypes by construction, and so does the immune context, which runs the other way: 2-hydroxyglutarate lowers STAT1 and CXCL10 and suppresses the accumulation of T cells in the tumour [15], and IDH-mutant gliomas express less PD-L1 than IDH-wildtype gliomas [16]. The two genotypes are therefore studied here as a biological contrast, and not as an arm from which a better response is expected.

Here we ask whether choline kinase α and PD-L1 regulate one another in GL261 glioma cells, and whether the IDH1 genotype alters that relation. We silenced each of the two genes and blocked PD-L1 with an antibody, and read the consequences by RT-qPCR, by RNA sequencing, by flow cytometry of surface PD-L1 and by high-resolution ^1^H NMR of dual-phase cell extracts. The two genes move in opposite directions in both genotypes, and that reciprocity reaches the protein at the cell surface in both. Blocking PD-L1 with an antibody does something the silencing does not, and what it does to choline metabolism and to lipid content stops at the IDH1 genotype.

## Results

### Silencing either gene raises the other, in both IDH1 genotypes

GL261 cells of IDH1-wildtype (GL261-WT) and IDH1-mutant (GL261-mIDH1) genotype were transfected with siRNA against *Chka*, against *Cd274*, or with a non-coding control, and transcripts were measured by RT-qPCR against that control.

Silencing *Chka* lowered its own transcript to ×0.51 (p = 0.034) in GL261-WT and to ×0.65 (p = 0.011) in GL261-mIDH1, and raised *Cd274* to ×3.08 (p = 0.047) and ×1.34 (p = 0.036) respectively (Fig. 1a). Silencing *Cd274* lowered it to ×0.50 (p = 0.065) and ×0.53 (p = 0.014), and raised *Chka* to ×1.78 (p = 0.0068) and ×1.28 (p = 0.015) (Fig. 1b). The two genes therefore moved in opposite directions whichever of them was silenced, and they did so in both genotypes.

**Figure 1.**
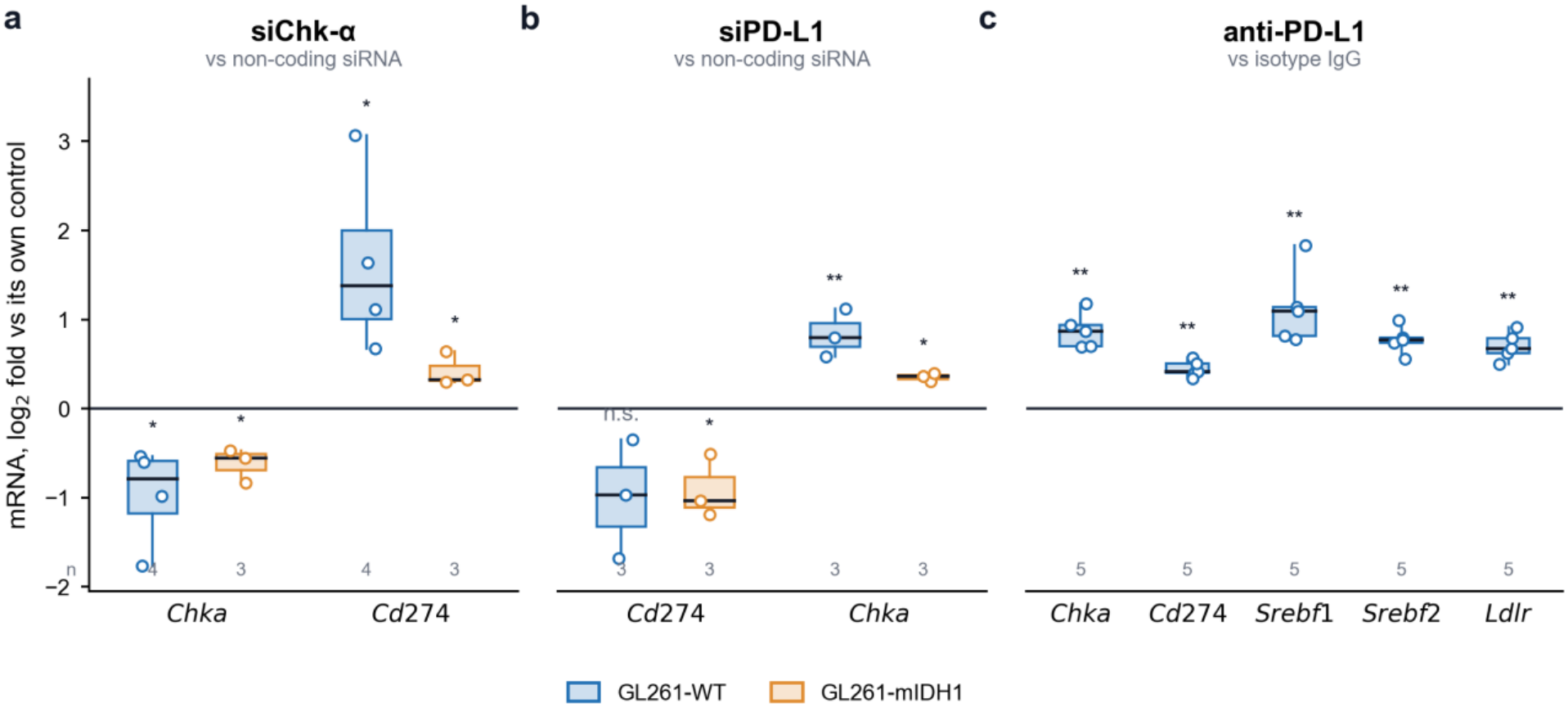
Silencing either gene raises the other, in both IDH1 genotypes, and PD-L1 blockade raises Chka together with the sterol regulators. RT-qPCR normalised to 18S in GL261-WT (blue) and GL261-mIDH1 (orange), 48 h after transfection or treatment, with each arm expressed against its own control. The y axis is the log_2_ fold change, so that zero is the control and one unit is a doubling. Boxes show the median and quartiles, whiskers the full range, and every independent experiment is shown as a point, with n given under each box. **(a)** siRNA against *Chka*, against non-coding siRNA. **(b)** siRNA against *Cd274*, against non-coding siRNA. **(c)** Anti-PD-L1 antibody, against isotype IgG, in GL261-WT only. Asterisks, unpaired two-tailed Student’s *t* test on log_2_ values against the control of each comparison, uncorrected for multiple testing: *p ≤ 0.05, **p < 0.01, ***p < 0.001; n.s., not significant.

RNA sequencing (RNA-seq) of the same perturbations reproduced both reciprocal contrasts. Silencing *Chka* raised *Cd274* ×2.3 in GL261-WT (adjusted p = 1.2 × 10^-5^) and ×2.6 in GL261-mIDH1 (adjusted p = 0.045), and silencing *Cd274* raised *Chka* ×1.4 in GL261-WT (adjusted p = 7.2 × 10^-4^) and ×1.3 in GL261-mIDH1, where it did not reach significance (adjusted p = 0.35). The knockdowns themselves were detected in GL261-WT, *Chka* falling to ×0.59 (adjusted p = 1.7 × 10^-10^) and *Cd274* to ×0.36 (adjusted p = 1.9 × 10^-6^) with their own siRNA. Neither knockdown was detected in the transcriptome of GL261-mIDH1, where both contrasts rested on fewer libraries.

### PD-L1 blockade raises *Chka* and the sterol regulators

Blocking PD-L1 with an antibody, on tumour cells cultured without immune cells, raised *Chka* to ×1.83 of the isotype IgG control (p = 0.0011) and *Cd274* to ×1.36 (p = 0.0086). In the same samples it raised the sterol regulators *Srebf1* to ×2.19 (p = 0.0044), *Srebf2* to ×1.71 (p = 0.0015) and *Ldlr* to ×1.62 (p = 0.0031) (Fig. 1c). By RNA sequencing, blockade raised *Chka* ×1.2 in GL261-WT, which did not reach significance (adjusted p = 0.14), and ×1.4 in GL261-mIDH1 (adjusted p = 0.028), and left *Cd274* unchanged in GL261-WT (×0.9; adjusted p = 0.81).

### Silencing ChoKα raises PD-L1 at the cell surface in both genotypes

Surface PD-L1 was measured by flow cytometry as the percentage of PD-L1-positive live cells (Fig. 2). Silencing *Chka* roughly doubled the PD-L1-positive fraction in both genotypes: ×2.12 in GL261-WT, from 6.6 ± 0.5 % in non-coding siRNA controls to 14.1 ± 3.6 % (p = 0.018), and ×1.95 in GL261-mIDH1, from 16.8 ± 7.5 % to 32.9 ± 1.0 % (p = 0.021). Against the vehicle control, which in GL261-mIDH1 sat lower and less variably at 5.1 ± 1.9 %, the same change was ×6.51 (p = 0.0002). Silencing *Cd274* lowered the positive fraction in GL261-WT to 2.1 ± 0.7 %, that is ×0.32 of the non-coding control (p = 0.0002) and ×0.27 of vehicle (p = 0.023). In GL261-mIDH1 it gave 8.4 ± 5.8 %, or ×0.50 of the non-coding control, which did not reach significance (p = 0.15).

**Figure 2.**
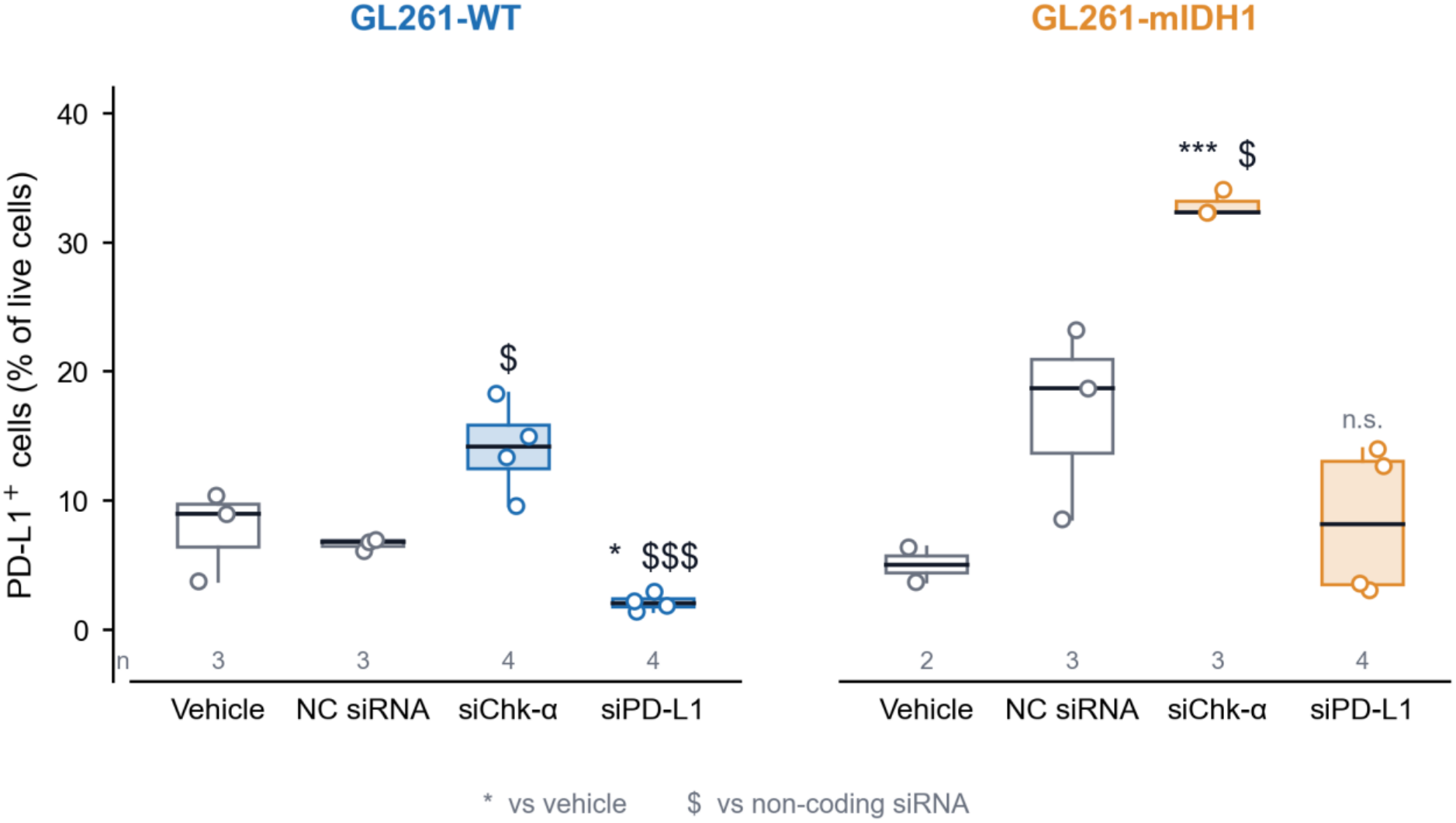
Silencing Chk-α raises surface PD-L1 in both genotypes and silencing PD-L1 lowers it in IDH1-wildtype cells. Percentage of PD-L1-positive cells among live cells in GL261-WT and GL261-mIDH1, measured by flow cytometry 48 h after transfection with non-coding siRNA, siRNA against *Chka* or siRNA against *Cd274*; vehicle denotes transfection reagent without siRNA. Boxes show the median and quartiles and whiskers the full range, with every independent experiment shown as a point and n given under each box. Asterisks, unpaired two-tailed Student’s *t* test against the vehicle control; dollar signs, the same test against the non-coding siRNA control: */$ p ≤ 0.05, **/$$ p < 0.01, ***/$$$ p < 0.001; n.s., not significant.

### The two siRNAs move phosphocholine in opposite directions, and the antibody lowers it

Dual-phase extracts of cells were analysed by high-resolution ^1^H NMR. Four resonances were resolved in the choline region, at 3.212, 3.222, 3.231 and 3.240 ppm, allowing phosphocholine to be quantified separately from free choline, glycerophosphocholine and the trimethylamine envelope (Fig. 3a).

**Figure 3.**
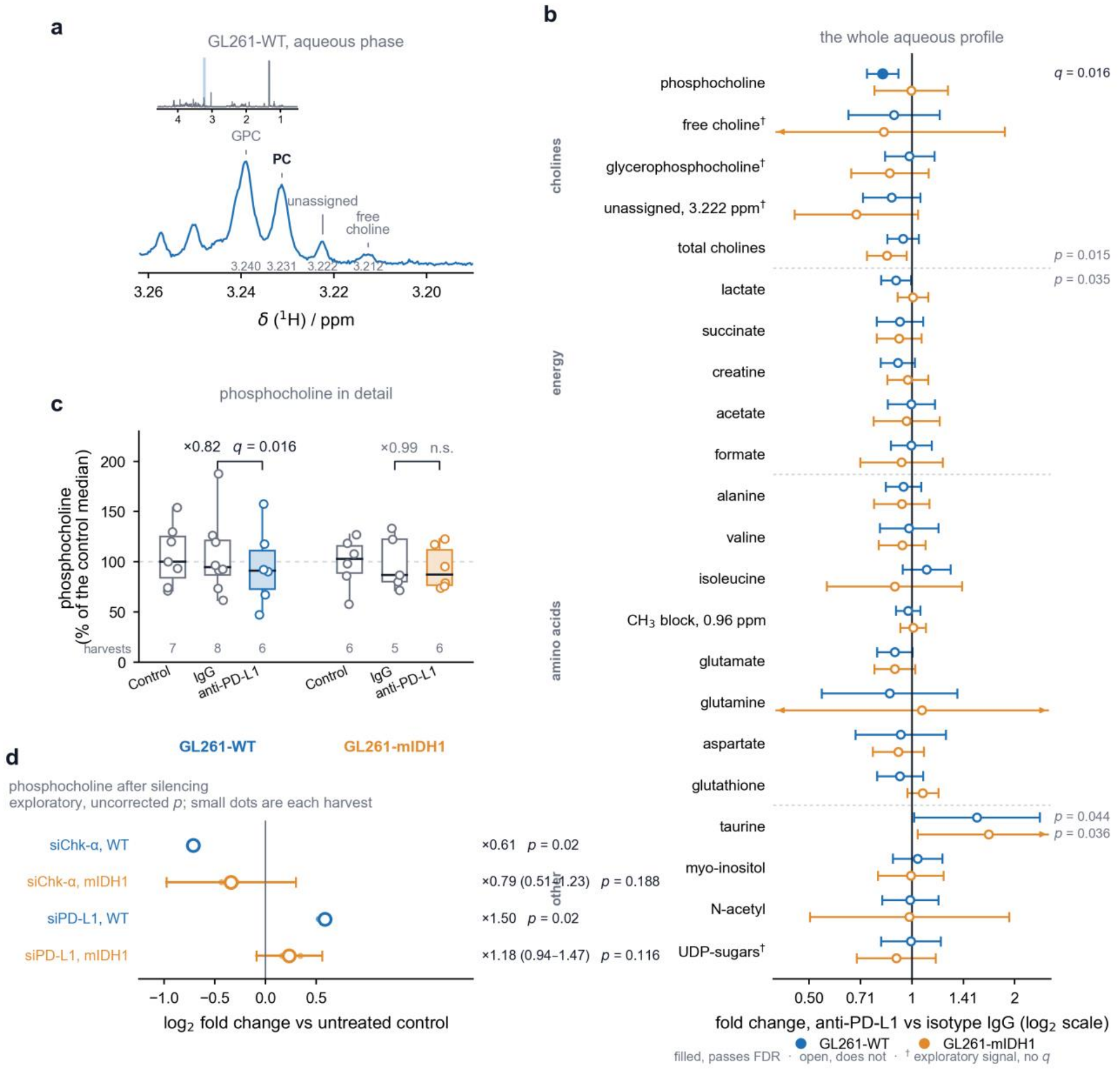
In the aqueous phase, phosphocholine is the only signal that responds to PD-L1 blockade, and the two siRNAs move it in opposite directions. **(a)** Choline region of a representative ^1^H NMR spectrum of an aqueous-phase extract of GL261-WT cells, with the four resolved resonances at 3.212 (free choline), 3.222 (unassigned), 3.231 (phosphocholine, PC) and 3.240 ppm (glycerophosphocholine, GPC). Inset, the same spectrum between 0.5 and 4.6 ppm, with the expanded region shaded. **(b)** The whole aqueous profile: fold change of each of the twenty-two resolved signals under antibody blockade against isotype IgG, in GL261-WT (blue) and GL261-mIDH1 (orange), with 95 % confidence intervals on a log_2_ scale; arrowheads mark intervals that run off the axis. Filled symbols pass the Benjamini–Hochberg correction applied across the panel within each genotype, open symbols do not, and † marks exploratory signals, which carry no q value. The q value is given for the signal that passes, and the uncorrected p for those below 0.05 that do not. **(c)** Phosphocholine in detail, profile-normalised, referred to cell number and expressed as a percentage of the median of the untreated control of its own genotype. Each point is one independent experiment, the mean of its tubes, and the number of experiments is given under each box; the fold change and its q value are given above each treated-against-isotype pair. **(d)** The siRNA arms, which are exploratory: fold change against untreated cells with its uncorrected p value, the small dots being the ratio of each experiment. Horizontal bars are 95 % confidence intervals where the design allows one; the GL261-WT arms rest on a single control per experiment and carry no interval.

In GL261-WT, silencing *Chka* lowered soluble phosphocholine to ×0.61 of untreated cells (p = 0.02), while silencing *Cd274* raised it to ×1.50 (p = 0.02). In GL261-mIDH1 both arms moved in the same directions without reaching significance, ×0.79 (p = 0.19) and ×1.18 (p = 0.12) (Fig. 3d). Antibody blockade lowered phosphocholine in the same direction as silencing *Chka*, and in the opposite direction to silencing *Cd274*. In GL261-WT it fell to ×0.82 of the isotype IgG control (95 % CI 0.74–0.91; p = 0.0009; q = 0.016), with the effect present in all six independent experiments. In GL261-mIDH1 there was no change (×0.99; p = 0.96) (Fig. 3c).

Across the whole aqueous profile, phosphocholine in GL261-WT was the only one of twenty-two signals whose response to blockade survived correction for multiple testing (Fig. 3b). Three further signals reached statistical significance: taurine rose in both genotypes, to ×1.55 in GL261-WT (p = 0.044) and ×1.68 in GL261-mIDH1 (p = 0.036); lactate fell to ×0.90 in GL261-WT (p = 0.035); and the total choline envelope fell to ×0.84 in GL261-mIDH1 (p = 0.015).

### PD-L1 blockade raises lipid content per cell in wild-type cells only

In the lipid phase of the same extracts (Fig. 4a), silencing *Cd274* raised the total lipid signal per cell in GL261-WT to ×1.10 of untreated cells without reaching significance (p = 0.14), a comparison that rests on a single experiment. Antibody blockade raised the same quantity to ×1.21 of the isotype IgG control in GL261-WT (p = 0.028; q = 0.081), with the effect present in all five experiments. In GL261-mIDH1 the same treatment gave ×0.84 (p = 0.46) (Fig. 4b). Tested as an interaction, the difference between the two genotypes was ×1.44 against untreated control cells (95 % CI 1.02–2.03; p = 0.039) and ×1.26 against the isotype IgG control (p = 0.31). Lipid composition did not change in either genotype: none of the thirteen resolved signals of the organic phase, cholesterol C18 among them, differed between treated and control cells (Fig. 4c).

**Figure 4.**
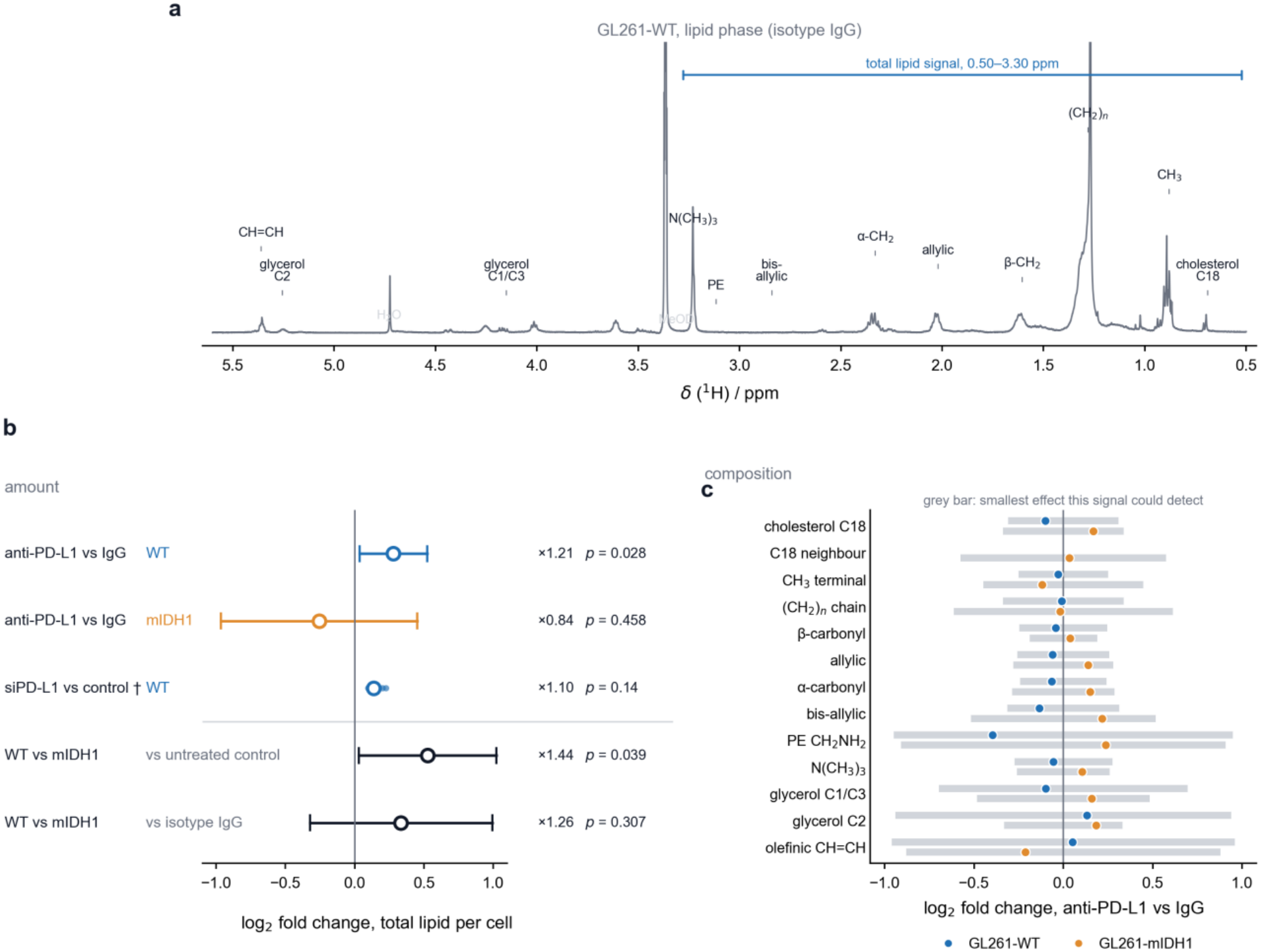
PD-L1 blockade raises lipid content per cell in IDH1-wildtype cells without changing lipid composition. **(a)** Representative ^1^H NMR spectrum of the organic phase of a dual-phase extract of GL261-WT cells treated with isotype IgG, with the resolved signals labelled and the 0.50–3.30 ppm window used for the total lipid signal marked. H_2_O and MeOD mark solvent resonances. **(b)** Total lipid signal per cell as the log_2_ fold change against the control of each comparison, with 95 % confidence intervals. † The siPD-L1 comparison rests on four tubes against a single control from one experiment and did not reach significance; its four tube ratios are shown as small dots. The difference between genotypes was tested in a linear model with the experiment as a block and is given against both controls, since the two do not agree. **(c)** Relative intensity of the resolved lipid signals, treated against isotype IgG; none differed in either genotype. The grey bar behind each point is the smallest effect **that signal** could have detected at 80 % power, which ranges from ×1.14 to ×1.95 and is therefore given per signal rather than as a single band. Total lipid signal per tube did not change; the increase in (b) appears when the signal is referred to cell number.

### The interferon response is engaged by every perturbation; the metabolic programmes are not

Gene-set enrichment analysis was performed on each of the three perturbations in each genotype (Fig. 5).

**Figure 5.**
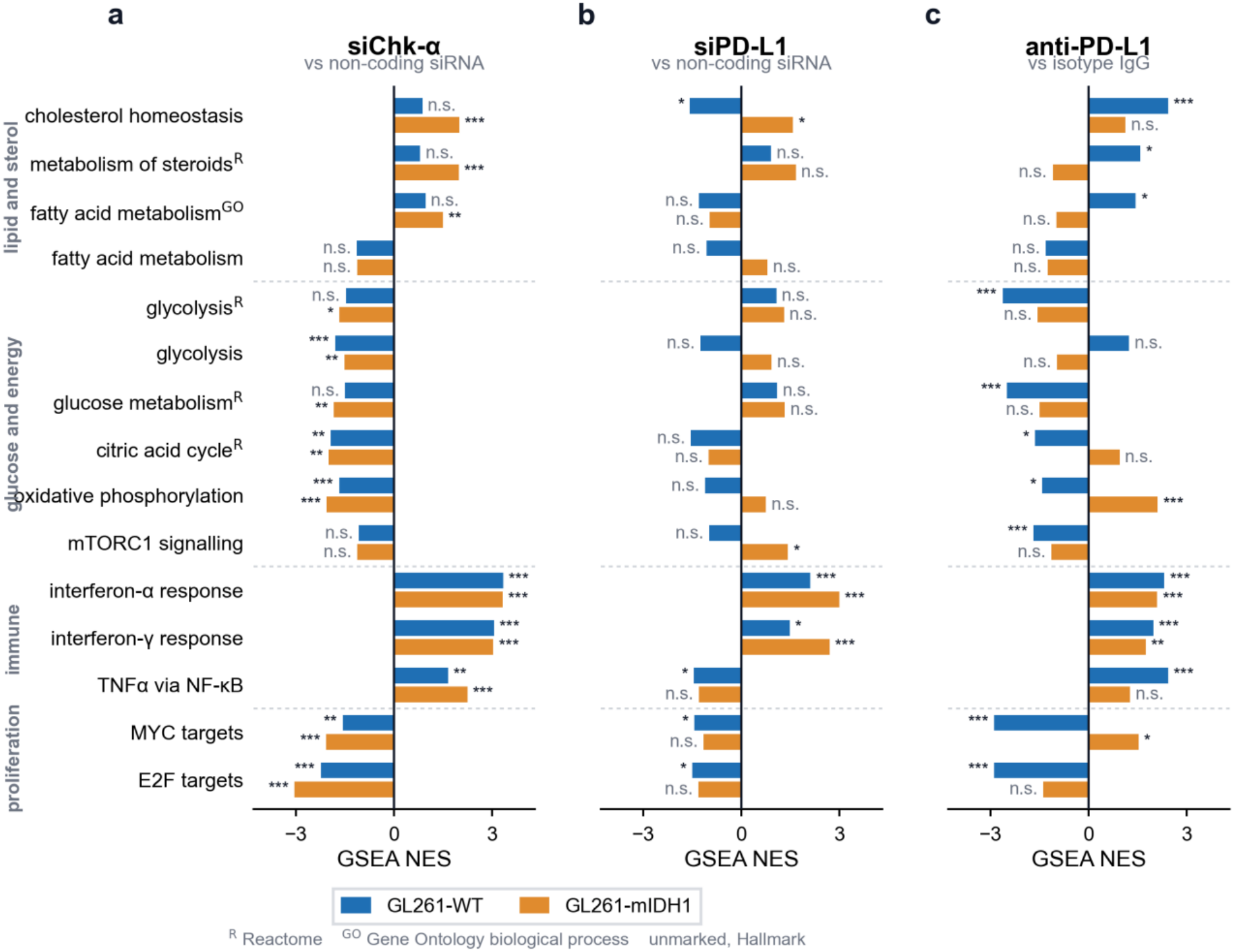
The interferon response is engaged by all three perturbations in both genotypes; the metabolic programmes are not. Normalised enrichment scores of fifteen gene sets under each of the three perturbations, in GL261-WT (blue) and GL261-mIDH1 (orange). **(a)** siRNA against *Chka*, against non-coding siRNA. **(b)** siRNA against *Cd274*, against non-coding siRNA. **(c)** Anti-PD-L1 antibody, against isotype IgG. Sets are grouped by the programme they belong to, and the collection each one comes from is marked in superscript: R, Reactome; GO, Gene Ontology biological process; unmarked, Hallmark. Where a Hallmark set covers the same process as a set from another collection, both are shown, so that no set is chosen for its result. Asterisks, Benjamini–Hochberg adjusted p value: *p ≤ 0.05, **p < 0.01, ***p < 0.001; n.s., not significant.

Silencing *Chka* enriched the interferon-α and interferon-γ response sets in both genotypes, with normalised enrichment scores of +3.34 and +3.06 in GL261-WT and +3.32 and +3.03 in GL261-mIDH1 (all adjusted p < 10^-22^), and depleted oxidative phosphorylation (−1.68 and −2.06), the citric acid cycle (−1.94 and −2.00), glycolysis (−1.81 and −1.53) and the MYC and E2F target sets in both. The sterol sets separated the two genotypes: cholesterol homeostasis and metabolism of steroids were enriched in GL261-mIDH1 (+1.99, adjusted p = 1.6 × 10^-4^; and +1.98, adjusted p = 3.8 × 10^-4^) and did not move in GL261-WT (+0.87 and +0.80, both not significant). In the same contrast *Stub1*, which destabilises PD-L1 protein, fell to ×0.42 in GL261-WT (adjusted p = 4.3 × 10^-19^) and ×0.53 in GL261-mIDH1 (adjusted p = 1.6 × 10^-4^), while *Cmtm6*, which stabilises it, rose to ×1.36 in GL261-WT (adjusted p = 1.2 × 10^-4^) and ×1.27 in GL261-mIDH1 (adjusted p = 0.35).

Silencing *Cd274* also enriched both interferon sets in both genotypes (+2.11 and +1.48 in GL261-WT, +2.99 and +2.69 in GL261-mIDH1), and moved little else. Cholesterol homeostasis was depleted in GL261-WT (−1.58, adjusted p = 0.040) and enriched in GL261-mIDH1 (+1.58, adjusted p = 0.042).

Antibody blockade enriched the interferon sets in both genotypes as the two siRNAs had done (+2.31 and +1.98 in GL261-WT, +2.09 and +1.74 in GL261-mIDH1), and in GL261-WT it was the perturbation that engaged the sterol programme most strongly: cholesterol homeostasis +2.43 (adjusted p = 6.3 × 10^-8^), metabolism of steroids +1.58 and the fatty acid metabolic process of Gene Ontology +1.44. In the same cells it depleted glycolysis (−2.63, adjusted p = 1.3 × 10^-10^), glucose metabolism (−2.50), the citric acid cycle (−1.65), oxidative phosphorylation (−1.42) and mTORC1 signalling (−1.68). The Hallmark sets covering glycolysis and fatty acid metabolism, shown alongside, did not move (+1.23 and −1.32).

In GL261-mIDH1 the same treatment left cholesterol homeostasis unchanged (+1.12, adjusted p = 0.46) and moved oxidative phosphorylation in the opposite direction (+2.11, adjusted p = 1.7 × 10^-7^). MYC targets were depleted in GL261-WT (−2.89) and enriched in GL261-mIDH1 (+1.52, adjusted p = 0.016).

## Discussion

An inverse dependence between choline kinase α and PD-L1 was first described in human triple-negative breast and pancreatic cancer cell lines, where silencing either gene raised the other and the relation was associated with COX-2 and TGF-β signalling [11]. The present data reproduce that relation in murine glioma cells, in two IDH1 genotypes, and add to it an antibody arm that the earlier work did not include and a metabolic response that stops at the IDH1 genotype.

Six observations of this work can be set against that one, and all six ran in the same direction (Table 1); five of them reached significance here. Silencing *Chka* raised *Cd274*, and silencing *Cd274* raised *Chka*; silencing *Chka* raised PD-L1 at the cell surface and lowered phosphocholine, and silencing *Cd274* raised phosphocholine. Only the rise in total lipid after PD-L1 silencing fell short of significance, and it moved in the published direction. The replication was obtained in another species, in another tumour, against another reference gene and in another laboratory. At the cell surface it reached both genotypes.

**Table 1.** The ChoKα–PD-L1 reciprocity described in human triple-negative breast and pancreatic cancer cells, set against the present measurements in murine GL261 glioma cells. Each row is a perturbation and the reading it was followed with. The published values are those reported in [11]; the present values are given against the control of each comparison, as in Figs 1–4.

| **Observation** | **Human TNBC / PDAC cells (*Cancer Metab* 2021) [11]** | **Murine GL261 cells (this work)** |
|---|---|---|
| **siChkα → PD-L1 transcript** | ↑ 85 ± 11 % (MDA-MB-231), 80 ± 15 % (SUM-149), 70 ± 10 % (Pa09C) | *Cd274* ×3.08 (p = 0.047) in WT and ×1.34 (p = 0.036) in mIDH1 |
| **siPD-L1 → Chk-α transcript** | ↑ > 50 % in both breast lines | *Chka* ×1.78 (p = 0.0068) in WT and ×1.28 (p = 0.015) in mIDH1 |
| **siChkα → surface PD-L1** | ↑ small but significant (% positive cells and MFI) | ×2.12 (p = 0.018) in WT and ×1.95 (p = 0.021) in mIDH1, against non-coding siRNA |
| **siChkα → phosphocholine** | ↓ significant | ×0.61 (p = 0.02) in WT |
| **siPD-L1 → phosphocholine** | ↑ significant | ×1.50 (p = 0.02) in WT |
| **siPD-L1 → total lipid** | ↑ | ×1.10 per cell (p = 0.14): same direction, not significant |

Blocking PD-L1 with an antibody and removing it with an siRNA did not do the same thing to phosphocholine. In IDH1-wildtype cells blockade lowered it to ×0.82 of the isotype control, in a planned and corrected comparison, while silencing raised it to ×1.50 of untreated cells. Silencing *Chka* lowered it to ×0.61, so the antibody moved the metabolite the way the choline kinase siRNA did and the opposite way to the PD-L1 siRNA. The two manoeuvres are not interchangeable: an antibody occupies the extracellular domain of a protein that stays where it is, whereas an siRNA removes the protein altogether, with its intracellular domain and whatever that domain does. PD-L1 does signal inside the tumour cell that carries it, since blocking it on tumour cells lowers their mTOR activity and their glycolysis [9], and attenuating it changes their growth and their autophagy in culture [10]. Our data place choline metabolism among the readings that separate the two manoeuvres; they do not identify what separates them.

That the metabolite in question is phosphocholine is not incidental. Of the twenty-two aqueous signals resolved in these extracts, it was the only one whose response to blockade survived correction for multiple testing, so the effect is specific rather than a general shift of the profile. It is also the branch of the lipid programme whose state is already read in the patient, because the total choline resonance of magnetic resonance spectroscopy follows the turnover of the phosphatidylcholine membrane that the Kennedy pathway builds. The four resonances resolved here measure phosphocholine apart from free choline and glycerophosphocholine, which the composite signal of an in vivo spectrum does not, and the fall measured here was a sixth of the pool, so whether a change of that size would be visible in vivo is not established by these data.

Both metabolic consequences of blockade were confined to the wild-type genotype: phosphocholine fell and lipid per cell rose in GL261-WT, and neither moved in GL261-mIDH1. The transcriptome drew the same line and in one place crossed it, with cholesterol homeostasis enriched in wild-type cells only and oxidative phosphorylation moving in opposite directions in the two genotypes. The interferon response did not follow that line. It was enriched by all three perturbations in both genotypes, most strongly of all by silencing *Chka*, and it is the only programme the three share; what the genotype bounds here is therefore the metabolic part of the response and not the inflammatory one. IDH1-mutant gliomas express less PD-L1 and carry a more methylated PD-L1 promoter than IDH1-wildtype gliomas [16], which would leave a smaller surface pool for an antibody to engage; that is one possible explanation and it is not tested here.

The sterol arm of the response to blockade was read twice in wild-type cells, in separate experiments. Gene-set enrichment placed cholesterol homeostasis at the head of the enriched sets, and RT-qPCR of the same treatment raised *Srebf1, Srebf2* and *Ldlr*, that is, both branches of the SREBP programme together with the receptor that imports cholesterol. Both readings are of transcript, so they agree without being independent of one another in kind. In the lipid phase of cells of the same genotype the signal per cell rose while the composition did not change, which makes the finding more lipid per cell and not a different lipid. The cholesterol resonance is one of the thirteen whose relative intensity did not move, and the smallest change it could have detected was ×1.24, so the sterol programme read in the transcriptome has no counterpart of that size in the composition of the lipid phase; nothing in these data links the rise in lipid to the fall in phosphocholine. Glioblastoma cells depend more on imported than on synthesised cholesterol [17], so a sterol programme that carries *Ldlr* with it would be compatible with uptake as much as with synthesis.

This study has limitations. The groups are small. The two transcript measurements did not agree on *Cd274* under blockade, which RT-qPCR raised and RNA sequencing did not, and disagreement of that kind between the two methods is documented [18]. Neither knockdown was detected in the transcriptome of the mutant line, where the contrasts rested on fewer libraries, so the transcriptomic arm of that genotype carries less weight than its wild-type counterpart. PD-L1 protein was measured at the cell surface and nowhere else, and only in the silencing arms: the blocking antibody and the detection antibody are the same clone, so surface PD-L1 read under blockade would report how much of the epitope the blocking antibody had left free rather than how much protein the cell carries. The surface arm therefore says nothing about what blockade does to the protein. All of this was done on tumour cells in culture, with no immune cells present.

If the reciprocity holds outside culture, lowering choline kinase α would raise PD-L1 on glioma cells, which would argue for combining the two targets rather than using them one after the other, and the IDH1 genotype may set how far such a combination reaches.

## Methods

### Cell lines and culture

GL261 murine glioma cells of IDH1-wildtype genotype (GL261-WT) were obtained from the American Type Culture Collection. GL261 cells carrying the IDH1-R132H mutation (GL261-mIDH1) were a gift from Dr Pellegata. Both lines were authenticated at the genomics service of the Instituto de Investigaciones Biomédicas Sols-Morreale and tested negative for mycoplasma.

Cells were maintained at 37 °C in a humidified atmosphere of 5 % CO_2_. GL261-WT cells were grown in RPMI 1640 (Sigma-Aldrich) supplemented with 10 % heat-inactivated fetal bovine serum (HyClone, Cytiva) and 1 % penicillin/streptomycin (Gibco). GL261-mIDH1 cells were grown in Dulbecco’s Modified Eagle Medium (Sigma-Aldrich) supplemented with 20 % heat-inactivated fetal bovine serum, 1 % penicillin/streptomycin and 0.5 mg/mL geneticin (Gibco). Cells were passaged at approximately 80 % confluence using TrypLE Express (Gibco) and counted with a CellDrop automated counter (DeNovix); all per-cell quantities reported in this work are referred to that count.

### Gene silencing

Cells were seeded, allowed to attach for 24 h, and transfected in fresh medium with siRNA at a final concentration of 100 nM using Lipofectamine RNAiMAX (Invitrogen, Thermo Fisher) according to the manufacturer’s instructions. Cells were harvested 48 h after transfection. A non-coding siRNA (MISSION siRNA Universal Negative Control #1, Sigma-Aldrich) was used as the negative control, and transfection reagent without siRNA as the vehicle control. All siRNAs were obtained from Merck (Sigma-Aldrich). The sense sequences were 5′-CAUGCUGUUCCAGUGCUCC-3′ against *Chka*, which matches both the murine and the human transcript, and 5′-GGUCAACGCCACAGCGAAU-3′ against murine *Cd274*.

### Antibody blockade of PD-L1

Cells were treated for 48 h with 10 µg/mL of anti-PD-L1 antibody (clone 10F.9G2, rat IgG2b κ) or with the same concentration of isotype-matched IgG, in the absence of immune cells.

### RT-qPCR

Transcripts were quantified by RT-qPCR and normalised to 18S ribosomal RNA. Each treatment arm was expressed relative to the control of its own plate: non-coding siRNA for the silencing arms and isotype IgG for the antibody arm. The murine *Cd274* primers were 5′-TTCAGATCACAGACGTCAAGCTG-3′ (forward) and 5′-ATTCTCTGGTTGATTTTGCGGTA-3′ (reverse), which give a 127 bp product on the *Cd274* transcript (NM_021893) and were checked for specificity by Primer-BLAST against the mouse RefSeq mRNA collection.

### Flow cytometry

Cells were seeded at 2.5 × 10^5^ per well, allowed to attach for 24 h, and transfected as described above. After 48 h they were harvested with TrypLE Express (Gibco) and resuspended in FACS buffer (phosphate-buffered saline with 1 % bovine serum albumin and 1 mM EDTA) containing Fc Block (BioLegend, 1:200) and either PE/Cy7-conjugated anti-PD-L1 (clone 10.F9G2, 1:400, BioLegend 124314) or the corresponding isotype control (clone RTK4530, 1:400, BioLegend 400617). After 30 min in the dark, cells were centrifuged at 1500 rpm for 5 min and resuspended in FACS buffer.

Samples were acquired on a FACSCanto II cytometer (Becton Dickinson), collecting a minimum of 30 000 events per sample, with To-Pro-3 (1:1000, Invitrogen T-3605) as a viability marker, and analysed with BD FACSDiva. The reported quantity is the percentage of PD-L1-positive cells among live, To-Pro-3-negative cells, with the positivity threshold set on the isotype control acquired in the same run.

### RNA sequencing

Libraries were prepared from independent experiments and sequenced paired-end; some samples were sequenced across two lanes. Library chemistry was not uniform across the whole set: part of the libraries were stranded and part unstranded, so the stranded libraries of the mixed set were requantified as unstranded from the same reads with the same index, which left read counts unchanged and raised mapping rates from 63–79 % to 83–86 %. In the antibody arm all libraries were quantified under a single convention.

Transcripts were quantified with Salmon 1.10.3 [19] against a GENCODE vM36 mouse index and summarised to gene level with tximport. Genes were retained when they reached at least 10 counts in at least 3 samples; pseudogenes, mitochondrially encoded genes, genes with a base mean below 20 and genes with an absolute log_2_ fold change above 5 were excluded.

Differential expression was computed with DESeq2 [20] within each cell line separately. Both arms were modelled with a term for the experiment and a term for the treatment, with contrasts taken against the non-coding siRNA control in the silencing arm and against isotype IgG in the antibody arm; the antibody model also carried library strandedness and standardised mapping rate. Significance was assessed on the Benjamini–Hochberg adjusted p value at 0.05, with no fold-change threshold. Genes for which DESeq2 returns no adjusted p value as a result of independent filtering are reported as not available.

As sensitivity analyses, each contrast was recomputed against the pooled control group, blocking by experiment day, jointly across the two lines, and, for the antibody arm, restricted to the natively stranded libraries; gene-level agreement between the alternative designs was r = 0.92–0.97.

### Gene-set enrichment analysis

Pre-ranked gene-set enrichment analysis [21] was performed with fgsea on the DESeq2 Wald statistic, restricted to sets of between 10 and 500 genes. Collections were Hallmark [22], Reactome, Gene Ontology biological process, WikiPathways and BioCarta, retrieved for mouse through msigdbr, together with twelve manually curated immune sets and twenty manually curated metabolic panels, one of which covers the choline and Kennedy pathway. The collection of origin is stated for every set named in the text and marked on the figure. Enrichment was assessed on the Benjamini–Hochberg adjusted p value.

### Dual-phase extraction

Cell pellets of a comparable number of cells were sonicated and transferred to glass tubes with 2.5–3 mL of methanol. After gentle vortexing, 6 mL of chloroform was added along the inner wall of the tube, and the sample was sonicated for 2–3 min with 1 s pulse intervals on ice and left for 1–2 h. Two millilitres of water were added, the sample was mixed by sonication for a further 30 s, and phase separation proceeded overnight at 4 °C. Samples were centrifuged at 4500 rpm for 25 min at 4 °C, and the upper aqueous phase and the lower organic phase were recovered separately with a glass Pasteur pipette into clean glass tubes, leaving the protein and cell-debris interface undisturbed.

The aqueous phase was dried under a nitrogen stream until approximately half its volume remained and then lyophilised. The lipid phase was dried under a nitrogen stream and lyophilised for 30–45 min. Dried samples were sealed and stored frozen until analysis.

Aqueous extracts were reconstituted in 600 µL of phosphate buffer in D_2_O at pH 7.4 containing 0.25 mM TSP as chemical-shift and quantification reference, vortexed, centrifuged at 4500 g for 5 min, and 550 µL of the supernatant transferred to the NMR tube. Lipid extracts were reconstituted in 600 µL of deuterated chloroform and methanol mixed 2:1 and containing 0.05 % TMS.

### NMR acquisition

Spectra were acquired on a Bruker Avance Neo spectrometer operating at 500.13 MHz for ^1^H, fitted with a 5 mm PA TXI 500W2/S4 H-C/N-D-03 Z inverse triple-resonance probe with z-gradient, at a sample temperature of 277.0 K.

Both phases were acquired with the same two one-dimensional experiments. The quantitative experiment was a single-pulse sequence with continuous-wave presaturation of the residual solvent during the relaxation delay (Bruker *zgpr*): 128 scans preceded by 8 dummy scans, a 5 µs excitation pulse at 14.66 W, a spectral width of 10 000 Hz (19.99 ppm) sampled over 65 536 complex points, which gives an acquisition time of 3.28 s, a relaxation delay of 2 s and therefore a recycle time of 5.28 s, a receiver gain of 101 and a pre-scan delay of 9.3 µs. The carrier was placed on the residual water resonance at 4.73 ppm, and the presaturation power was set per sample and recorded for use as a covariate (see below), its typical values being 4 × 10^-4^ W in aqueous extracts and 3 × 10^-5^ W in lipid extracts. These are pulse-acquire experiments and carry no echo time. A survey spectrum without presaturation was also acquired for every sample with a 30° excitation pulse (Bruker *zg30*), 4 scans and a relaxation delay of 1 s; in the lipid phase the olefinic resonance, which lies under the suppressed region of the presaturation spectrum, was measured on this survey spectrum, and the remaining twelve lipid signals on the presaturation spectrum.

Aqueous spectra were referenced to TSP at 0.000 ppm and lipid spectra to the chain methylene resonance at 1.27 ppm.

### NMR processing and quantification

Spectra were processed from the free induction decay with a single procedure comprising 0.3 Hz exponential line broadening, zero-filling to 65 536 points, digital-filter correction, automatic phasing subsequently inspected, chemical-shift referencing to TSP or to the chain methylene, and interpolation onto a common grid from −0.5 to 10.0 ppm in steps of 0.0005 ppm. Intensities were scaled as area × 2^NC × 128/NS so that areas are comparable between spectra. Baselines were corrected with a Whittaker smoother in the aqueous phase and with a second-order polynomial fitted in signal-free windows in the lipid phase.

Because presaturation attenuates resonances close to the solvent signal in proportion to the applied power, and that power varied between acquisitions, each area was multiplied by F(δ) = 1 + k_1_·PLW9/(O1 − δ)^2^, where δ is the position of the signal, O1 that of the carrier and PLW9 the presaturation power. The constant k_1_ was calibrated using samples measured twice at different powers, giving k_1_ = 684 for aqueous extracts; lipid spectra were calibrated separately. All principal results were recomputed across k_1_ from 0 to 2000 and are unchanged.

Signal lists were fixed before any statistical test was computed and without reference to treatment identity: 18 result signals in the aqueous phase and 26 in the lipid phase, of which 12 were analysed in GL261-WT and 13 in GL261-mIDH1. In the choline region, four resonances were fitted separately: 3.212 ppm (free choline), 3.222 ppm (unassigned), 3.231 ppm (phosphocholine) and 3.240 ppm (glycerophosphocholine). In the lipid phase, total lipid is the integral between 0.50 and 3.30 ppm.

Composition was assessed on probabilistic quotient normalised spectra, computed over 0.50–3.30 ppm in the lipid phase. Quantities per cell are absolute areas divided by the cell count of the same culture. Every composition result was also examined under at least one other normalisation.

### Statistical analysis

RT-qPCR and flow cytometry data were compared with unpaired two-tailed Student’s *t* tests, on log_2_-transformed ratios for RT-qPCR and on percentages for flow cytometry, against the control indicated in each case. These p values are not corrected for multiple testing. Flow cytometry data are given as mean ± standard deviation and each replicate is an independent experiment.

NMR comparisons specified in advance were analysed with a linear model on log_2_-transformed values of the form log_2_(value) ∼ treatment + experiment (block) + covariate, the covariate being presaturation power for aqueous extracts and the line width of the cholesterol C18 resonance for lipid extracts. The principal analysis compares the two groups of interest, the isotype IgG being the control for antibody treatment; a three-group model including untreated cells was fitted alongside. P values were corrected for multiple testing by the Benjamini–Hochberg procedure within each comparison and cell line, across the result signals, and are reported as q values. Robustness was assessed by removing one sample at a time, removing one experiment at a time, varying the model, sweeping the presaturation constant, permuting treatment labels within experiment 2000 times, and computing the minimum effect detectable at 80 % power, which is reported for every signal of the lipid phase.

Analyses not specified in advance are identified as exploratory and their p values are reported uncorrected. These comprise least-squares models on log_2_ values with treatment and experiment as a block, fitted only on experiments containing both groups; Welch *t* tests and Mann–Whitney tests; the Crawford–Howell single-case test where a treated group is compared against a single control; and, for the direct comparison between the two genotypes, a single model fitted across both lines with line × experiment as the block and a treatment × line interaction term. The two genotypes are not otherwise compared with each other.

## Data availability

RNA sequencing data and NMR spectra generated in this study are being deposited in the Gene Expression Omnibus and in MetaboLights respectively. Accession numbers will be added to a revised version of this preprint and will be public on or before journal publication.

## Notes

### Competing Interest Statement

The authors have declared no competing interest.

### Funding

- Agencia Estatal de Investigación: PID2022-137572OA-I00

## References

[1] Stupp R, Hegi ME, Mason WP, van den Bent MJ, Taphoorn MJ, Janzer RC, Ludwin SK, Allgeier A, Fisher B, Belanger K, Hau P, Brandes AA, Gijtenbeek J, Marosi C, Vecht CJ, Mokhtari K, Wesseling P, Villa S, Eisenhauer E, Gorlia T, Weller M, Lacombe D, Cairncross JG, Mirimanoff RO. Effects of radiotherapy with concomitant and adjuvant temozolomide versus radiotherapy alone on survival in glioblastoma in a randomised phase III study: 5-year analysis of the EORTC-NCIC trial. The Lancet Oncology 2009;10(5):459–466. DOI: 10.1016/S1470-2045(09)70025-7 · PMID: 19269895

[2] Omuro A, Brandes AA, Carpentier AF, Idbaih A, Reardon DA, Cloughesy T, Sumrall A, Baehring J, van den Bent M, Bähr O, Lombardi G, Mulholland P, Tabatabai G, Lassen U, Sepulveda JM, Khasraw M, Vauleon E, Muragaki Y, Di Giacomo AM, Butowski N, Roth P, Qian X, Fu AZ, Liu Y, Potter V, Chalamandaris AG, Tatsuoka K, Lim M, Weller M. Radiotherapy combined with nivolumab or temozolomide for newly diagnosed glioblastoma with unmethylated MGMT promoter: An international randomized phase III trial. Neuro-Oncology 2023;25(1):123–134. DOI: 10.1093/neuonc/noac099 · PMID: 35419607

[3] Reardon DA, Brandes AA, Omuro A, Mulholland P, Lim M, Wick A, Baehring J, Ahluwalia MS, Roth P, Bähr O, Phuphanich S, Sepulveda JM, De Souza P, Sahebjam S, Carleton M, Tatsuoka K, Taitt C, Zwirtes R, Sampson J, Weller M. Effect of Nivolumab vs Bevacizumab in Patients With Recurrent Glioblastoma: The CheckMate 143 Phase 3 Randomized Clinical Trial. JAMA Oncology 2020;6(7):1003–1010. DOI: 10.1001/jamaoncol.2020.1024 · PMID: 32437507

[4] Cloughesy TF, Mochizuki AY, Orpilla JR, Hugo W, Lee AH, Davidson TB, Wang AC, Ellingson BM, Rytlewski JA, Sanders CM, Kawaguchi ES, D. L, Li G, Yong WH, Gaffey SC, Cohen AL, Mellinghoff IK, Lee EQ, Reardon DA, O’Brien BJ, Butowski NA, Nghiemphu PL, Clarke JL, Arrillaga-Romany IC, Colman H, Kaley TJ, de Groot JF, Liau LM, Wen PY, Prins RM. Neoadjuvant anti-PD-1 immunotherapy promotes a survival benefit with intratumoral and systemic immune responses in recurrent glioblastoma. Nature Medicine 2019;25(3):477–486. DOI: 10.1038/s41591-018-0337-7 · PMID: 30742122

[5] Glunde K, Bhujwalla ZM, Ronen SM. Choline metabolism in malignant transformation. Nature Reviews Cancer 2011;11(12):835–848. DOI: 10.1038/nrc3162 · PMID: 22089420

[6] Ramírez de Molina A, Rodríguez-González A, Gutiérrez R, Martínez-Piñeiro L, Sánchez J, Bonilla F, Rosell R, Lacal J. Overexpression of choline kinase is a frequent feature in human tumor-derived cell lines and in lung, prostate, and colorectal human cancers. Biochemical and Biophysical Research Communications 2002;296(3):580–583. DOI: 10.1016/s0006-291x(02)00920-8 · PMID: 12176020

[7] Venkatesh HS, Chaumeil MM, Ward CS, Haas-Kogan DA, James CD, Ronen SM. Reduced phosphocholine and hyperpolarized lactate provide magnetic resonance biomarkers of PI3K/Akt/mTOR inhibition in glioblastoma. Neuro-Oncology 2011;14(3):315–325. DOI: 10.1093/neuonc/nor209 · PMID: 22156546

[8] Koch K, Hartmann R, Schröter F, Suwala AK, Maciaczyk D, Krüger AC, Willbold D, Kahlert UD, Maciaczyk J. Reciprocal regulation of the cholinic phenotype and epithelial-mesenchymal transition in glioblastoma cells. Oncotarget 2016;7(45):73414–73431. DOI: 10.18632/oncotarget.12337 · PMID: 27705917

[9] Chang CH, Qiu J, O’Sullivan D, Buck MD, Noguchi T, Curtis JD, Chen Q, Gindin M, Gubin MM, van der Windt GJW, Tonc E, Schreiber RD, Pearce EJ, Pearce EL. Metabolic Competition in the Tumor Microenvironment Is a Driver of Cancer Progression. Cell 2015;162(6):1229–1241. DOI: 10.1016/j.cell.2015.08.016 · PMID: 26321679

[10] Clark CA, Gupta HB, Sareddy G, Pandeswara S, Lao S, Yuan B, Drerup JM, Padron A, Conejo-Garcia J, Murthy K, Liu Y, Turk MJ, Thedieck K, Hurez V, Li R, Vadlamudi R, Curiel TJ. Tumor-Intrinsic PD-L1 Signals Regulate Cell Growth, Pathogenesis, and Autophagy in Ovarian Cancer and Melanoma. Cancer Research 2016;76(23):6964–6974. DOI: 10.1158/0008-5472.CAN-16-0258 · PMID: 27671674

[11] Pacheco-Torres J, Penet MF, Mironchik Y, Krishnamachary B, Bhujwalla ZM. The PD-L1 metabolic interactome intersects with choline metabolism and inflammation. Cancer & Metabolism 2021;9(1):10. DOI: 10.1186/s40170-021-00245-w · PMID: 33608051

[12] Louis DN, Perry A, Wesseling P, Brat DJ, Cree IA, Figarella-Branger D, Hawkins C, Ng HK, Pfister SM, Reifenberger G, Soffietti R, von Deimling A, Ellison DW. The 2021 WHO Classification of Tumors of the Central Nervous System: a summary. Neuro-Oncology 2021;23(8):1231–1251. DOI: 10.1093/neuonc/noab106 · PMID: 34185076

[13] Yan H, Parsons DW, Jin G, McLendon R, Rasheed BA, Yuan W, Kos I, Batinic-Haberle I, Jones S, Riggins GJ, Friedman H, Friedman A, Reardon D, Herndon J, Kinzler KW, Velculescu VE, Vogelstein B, Bigner DD. IDH1 and IDH2 mutations in gliomas. New England Journal of Medicine 2009;360(8):765–773. DOI: 10.1056/NEJMoa0808710 · PMID: 19228619

[14] Mellinghoff IK, van den Bent MJ, Blumenthal DT, Touat M, Peters KB, Clarke J, Mendez J, Yust-Katz S, Welsh L, Mason WP, Ducray F, Umemura Y, Nabors B, Holdhoff M, Hottinger AF, Arakawa Y, Sepulveda JM, Wick W, Soffietti R, Perry JR, Giglio P, de la Fuente M, Maher EA, Schoenfeld S, Zhao D, Pandya SS, Steelman L, Hassan I, Wen PY, Cloughesy TF. Vorasidenib in IDH1- or IDH2-Mutant Low-Grade Glioma. New England Journal of Medicine 2023;389(7):589–601. DOI: 10.1056/NEJMoa2304194 · PMID: 37272516

[15] Kohanbash G, Carrera DA, Shrivastav S, Ahn BJ, Jahan N, Mazor T, Chheda ZS, Downey KM, Watchmaker PB, Beppler C, Warta R, Amankulor NA, Herold-Mende C, Costello JF, Okada H. Isocitrate dehydrogenase mutations suppress STAT1 and CD8+ T cell accumulation in gliomas. The Journal of Clinical Investigation 2017;127(4):1425–1437. DOI: 10.1172/JCI90644 · PMID: 28319047

[16] Berghoff AS, Kiesel B, Widhalm G, Wilhelm D, Rajky O, Kurscheid S, Kresl P, Wöhrer A, Marosi C, Hegi ME, Preusser M. Correlation of immune phenotype with IDH mutation in diffuse glioma. Neuro-Oncology 2017;19(11):1460–1468. DOI: 10.1093/neuonc/nox054 · PMID: 28531337

[17] Villa GR, Hulce JJ, Zanca C, Bi J, Ikegami S, Cahill GL, Gu Y, Lum KM, Masui K, Yang H, Rong X, Hong C, Turner KM, Liu F, Hon GC, Jenkins D, Martini M, Armando AM, Quehenberger O, Cloughesy TF, Furnari FB, Cavenee WK, Tontonoz P, Gahman TC, Shiau AK, Cravatt BF, Mischel PS. An LXR-Cholesterol Axis Creates a Metabolic Co-Dependency for Brain Cancers. Cancer Cell 2016;30(5):683–693. DOI: 10.1016/j.ccell.2016.09.008 · PMID: 27746144

[18] Everaert C, Luypaert M, Maag JLV, Cheng QX, Dinger ME, Hellemans J, Mestdagh P. Benchmarking of RNA-sequencing analysis workflows using whole-transcriptome RT-qPCR expression data. Scientific Reports 2017;7(1):1559. DOI: 10.1038/s41598-017-01617-3 · PMID: 28484260

[19] Patro R, Duggal G, Love MI, Irizarry RA, Kingsford C. Salmon provides fast and bias-aware quantification of transcript expression. Nature Methods 2017;14(4):417–419. DOI: 10.1038/nmeth.4197 · PMID: 28263959

[20] Love MI, Huber W, Anders S. Moderated estimation of fold change and dispersion for RNA-seq data with DESeq2. Genome Biology 2014;15(12):550. DOI: 10.1186/s13059-014-0550-8 · PMID: 25516281

[21] Subramanian A, Tamayo P, Mootha VK, Mukherjee S, Ebert BL, Gillette MA, Paulovich A, Pomeroy SL, Golub TR, Lander ES, Mesirov JP. Gene set enrichment analysis: a knowledge-based approach for interpreting genome-wide expression profiles. Proceedings of the National Academy of Sciences of the United States of America 2005;102(43):15545–15550. DOI: 10.1073/pnas.0506580102 · PMID: 16199517

[22] Liberzon A, Birger C, Thorvaldsdóttir H, Ghandi M, Mesirov JP, Tamayo P. The Molecular Signatures Database (MSigDB) hallmark gene set collection. Cell Systems 2015;1(6):417–425. DOI: 10.1016/j.cels.2015.12.004 · PMID: 26771021

